# A comprehensive DNA methylation atlas for the Chinese population through nanopore long-read sequencing of 106 individuals

**DOI:** 10.64898/2026.04.20.719515

**Authors:** Yang Li, Tao Jiang, Long Qian, Yadong Wang

## Abstract

DNA methylation constitutes the primary epigenetic language mediating organismal phenotypic plasticity. Establishing a cohort-level genomic methylation landscape featuring wide geographical diversity is fundamental for dissecting its genetic and environmental attributes. Leveraging nanopore sequencing’s strength in genome-methylome co-sequencing, we generated a whole-genome, haplotype-resolved methylation atlas for 106 individuals from 19 provinces across China. The atlas identified 27,609,354 CpG sites genome-wide, with notably more informed gene proximal regions and CpG islands compared to whole-genome bisulfite sequencing. Detailed analyses revealed genomic structural variants as a pervasive covariate of DNA methylation, with a remarkable 2-fold compensation effect found in genome-wide heterozygous deletions. On the other hand, habitat altitude is found to be a strong environmental determinant of DNA methylation. We established a quantitative relationship between altitude and methylation states and identified a gene set strictly responsive to altitude differences, revealing epigenetically regulated genes such as *PRDM16, EPHB2* and *WNT7A*. The methylation atlas provides a reference resource to facilitate further explorations into human epigenetics.

## Introduction

DNA methylation, a pivotal epigenetic modification, is intricately influenced by both genetic and environmental factors and plays a crucial role in environmental adaptation. Among various epigenetic marks, 5-methylcytosine (5mC) is the most prevalent and functionally significant methylation modification in the human genome, exerting profound effects on gene transcriptional regulation^1, 2^. Studies using whole-genome bisulfite sequencing (WGBS) have laid a foundation for this epigenetic landscape. However, the bisulfite base conversion reaction may introduce biases; meanwhile, short-read sequencing has inherent limitations in resolving complex genomic regions^3, 4^. Recent advances in nanopore sequencing have addressed these challenges by overcoming read-length constraints and eliminating the need for chemical modifications that compromise DNA integrity^5, 6^. With the ability to generate reads spanning tens of thousands of bases, nanopore sequencing enhances the detection of complex structural variants (SVs) and enables highly contiguous haplotype phasing ^7, 8^. Furthermore, its capacity to directly sequence native DNA allows for the simultaneous identification of 5mC at single-nucleotide resolution^9^, facilitating the integration of genomic and epigenomic data in a single assay. These technological advancements offer unprecedented opportunities for in-depth explorations into genome-wide epigenetic mechanisms and the interplay between environment and epigenetic regulation.

Large-scale cohort studies have widely applied DNA methylation profiling to investigate its role in biological processes on multiple scales. For example, the construction of a comprehensive human cell DNA methylation atlas has revealed highly consistent methylation patterns within identical cell types^10, 11^. Similarly, the development of a large-scale DNA methylation QTL (meQTL) atlas has provided valuable insights into the impact of methylation in regulatory regions on human diseases^12^. More recently, research on population-specific DNA methylation has identified substantial epigenetic differences among human populations, shedding light on the relationship between epigenetic variation and environmental adaptability^13^. A study conducted across diverse African environments has shown that DNA methylation changes, particularly those associated with historical lifestyles, are shaped by natural selection^14^. As one of the most populous countries in the world, China has also conducted relevant studies, encompassing four populations including Tibetan and two populations including Uyghur to characterize DNA methylation patterns^15, 16^, providing valuable insights into population-specific epigenetic features. These findings underscore the critical role of DNA methylation in gene expression regulation, emphasizing its significance as a key mediator of genetic and environmental interactions.

While inter-population differences have been widely studied in these works, allele- and haplotype-specific methylation studies remain underrepresented in population-comparative settings^17, 18^. Likewise, the utilization of WGBS in these studies resulted in an incomplete representation of the methylation landscape, leaving out complex genomic regions. In particular, it limits interrogation on the interplay between genome-wide SVs and DNA methylation^19^. Furthermore, existing studies have contrasted samples from extreme environments (e.g., high-vs. low altitude, hot vs. cold climates), but the absence of data from transitional environments has hindered quantitative assessment of environmental influences on DNA methylation^20, 21, 22^. Therefore, there is an urgent need to establish a comprehensive methylation atlas that captures the dynamics of epigenetic modifications along continuous environmental gradients, enables high-resolution genome-wide methylation profiling, and delineates finer-scale methylation patterns at the allele level.

Herein, we constructed a comprehensive methylation atlas for 106 individuals from 19 provinces in China across an environmental gradient using whole-genome nanopore sequencing (Fig. 1). Based on this atlas, we identified three types of differentially methylated regions (DMRs)—segment-related DMRs (sDMRs), haplotype-based DMRs (hDMRs), and population-specific DMRs (pDMRs)—capturing 5mC profile differences at multiple scales including the genome level, the haplotype level and the population level. Our study reveals a striking two-fold compensatory methylation effect at heterozygous deletion sites, presumably to maintain gene dosage control, while mobile element insertions exhibit systematic hypomethylation, suggestive of genome-wide epigenetic processes. Furthermore, we demonstrate that high-altitude epigenetic adaptation operates on multiple scales, with haplotype-specific regulation at the individual level and progressive adaptation at the population level, collectively shaping a dynamic and adaptive epigenetic landscape. These analyses uncovered novel genes closely linked to high-altitude adaptation. The full methylation atlas is open-sourced on a user-friendly visualization website to facilitate future cutting-edge genomic and epigenomic studies.

**Figure 1.**
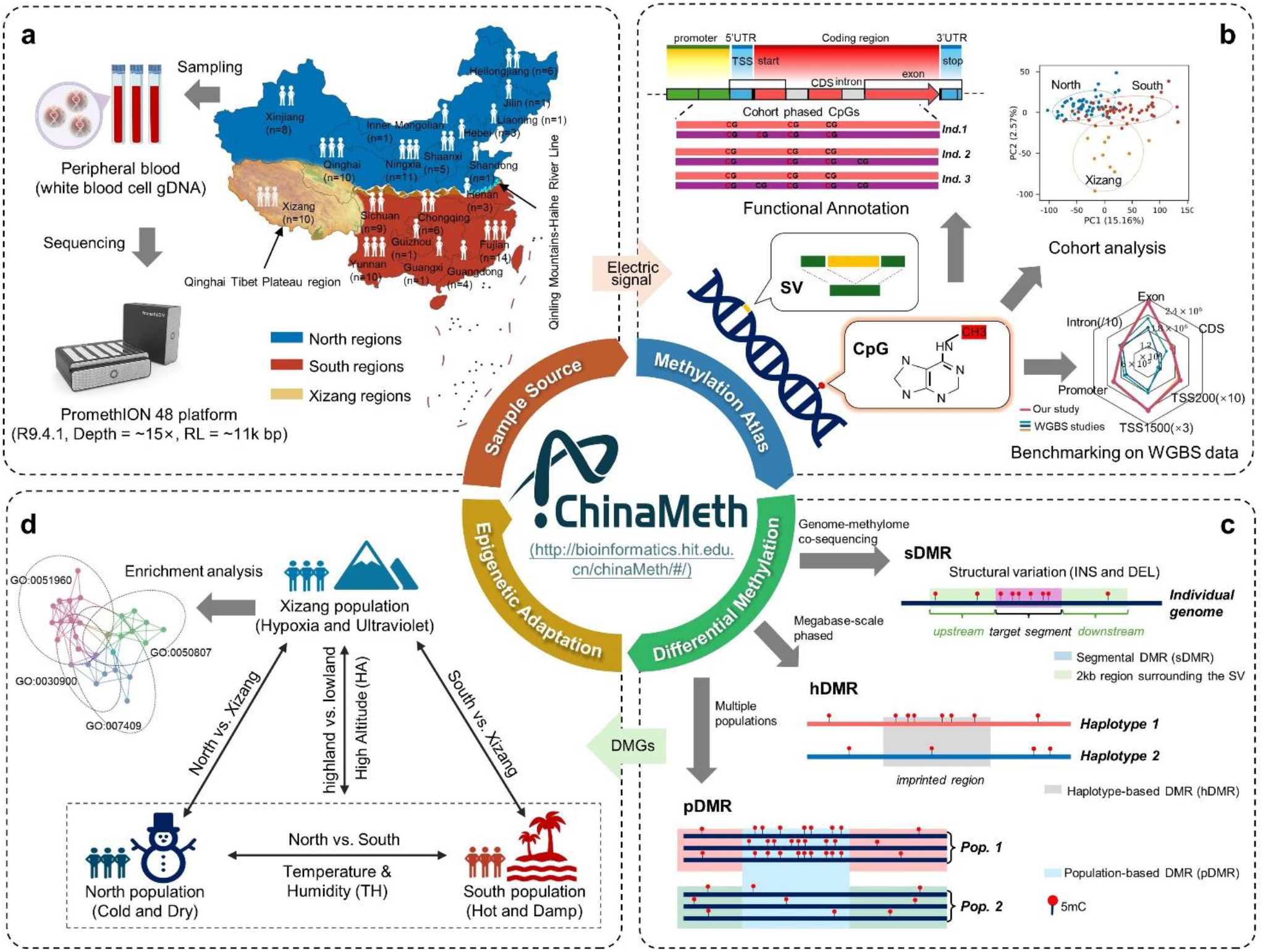
Overview of DNA methylation analysis in the ChinaMeth project. **(a)** Sample collection and sequencing. Peripheral blood samples were collected from individuals in North, South, and Xizang regions. Whole-genome nanopore sequencing was performed using PromethION (R9.4.1). **(b)** Methylation atlas construction. CpG methylation and SV signals were extracted after alignment to GRCh38. Single nucleotide variant (SNV)-based haplotype phasing was applied for allele-specific methylation analysis. Methylation profiles were annotated, compared across populations, and benchmarked against WGBS data. **(c)** DMR identification. sDMRs were identified by integrating methylation and insertion/deletion signals. hDMRs were derived from accurately phased haplotypes, allowing allele-specific resolution. pDMRs were identified by comparing methylation patterns across populations. **(d)** Epigenetic adaptation analysis. DMRs were used to define population-specific differentially methylated genes (DMGs) which were further modeled along defined environmental factors. The ChinaMeth web platform enables exploration and visualization of methylation patterns.

## Results

### Construction of a Chinese population methylation atlas by nanopore sequencing

To construct a comprehensive DNA methylation atlas representative of the Chinese population, we selected 106 healthy individuals from five biobanks with broad geographical distribution across China: 48 individuals from the North (10 provinces), 48 from the South (8 provinces), and 10 from Xizang (Fig. 2a). The sampled regions span altitudes from 50 to 5,337 meters, with average temperatures ranging from -5°C to 23.5°C and relative humidity varying between 37.2% and 80%, covering the broadest environmental landscapes in China. Genomic DNA was extracted following stringent quality control protocols and sequenced on the Oxford Nanopore Technologies (ONT) PromethION 48 platform with R9.4 flowcells and chemistry. Base calling was performed using Dorado^23^ (v0.3.1), achieving an average coverage of 14.6× (10–24×). The average sequencing error rate was **∼**6.8%, and the average read N50 was 14,179 bp. Following read and CpG site filtering (primary alignments with MAPQ ≥ 5 and CpG site coverage ≥ 3–8×), methylation detection and haplotype-phasing indicated that each individual contained an average of 26,276,583 CpGs (excluding sex chromosomes) with bimodally distributed methylation levels averaging at 80.74%. A more stringent mapping quality (MAPQ ≥ 20) produced nearly identical results (99% CpG overlap and *r* = 0.98 correlation of methylation levels). Therefore, to maximize genome-wide CpG coverage for comprehensive discovery, all analyses were done with the MAPQ ≥ 5 dataset. Additionally, an average of 19,636 SVs were identified across autosomal chromosomes. Down-sampling revealed that detected CpG counts plateaued at the current coverage level (Fig. 2b). Furthermore, no significant differences in CpG counts were observed among the three populations (Wilcoxon rank-sum tests, p > 0.05, Fig. 2b), confirming that the coverage range employed in this study is sufficient for genome-wide CpG analysis.

**Figure 2.**
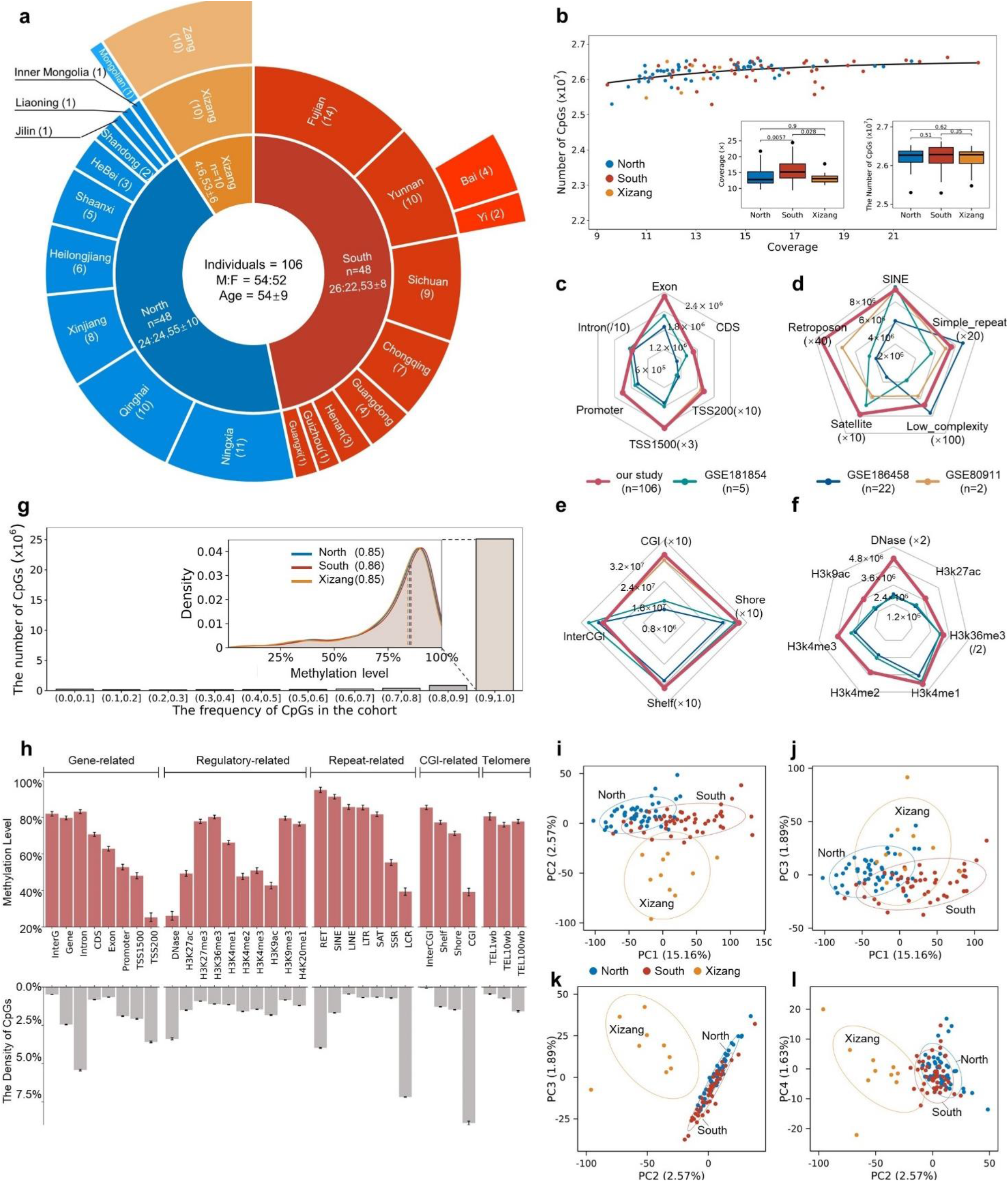
Comprehensive characterization of the methylation atlas. **(a)** Cohort composition showing 106 participants represented by concentric rings showing regional (inner circle), provincial (middle circle), and ethnic (outer circle) classifications. Bracketed numbers indicate sample sizes. Participants were recruited from North China, South China, and Xizang, with balanced sex ratios (Male: Female) and comparable age distributions (mean ± SD). **(b)** Relationship between sequencing coverage and detected CpG counts, with a loess regression curve demonstrating a positive trend. Inset boxplots display distributions of sequencing coverage (*x*-axis) and corresponding CpG counts (*y*-axis) across populations. **(c–f)** Comparison of detected CpGs by nanopore sequencing (this study) and in three WGBS datasets across genomic regions: gene-related **(c)**, repeat **(d)**, CpG island-associated **(e)**, and regulatory regions **(f)**. Dataset sample sizes are shown in the legend; specific axes are scaled with folds indicated in brackets. **(g)** Conservation of CpGs across individuals, revealing most CpGs (>90% cohort-wide) are shared. Inset: Kernel density estimation of methylation levels on CpGs with population frequencies in the (0.9, 1] range; dashed lines indicate median methylation in three populations. **(h)** Genome regional CpG density (lower panel) and methylation levels (upper panel). InterG: intergenic; RTP: retroposon; SAT: satellite; SSR: simple repeat; LCR: low complexity; TEL: telomere. **(i–l)** Principal component analysis of the first four components. Dashed ellipses indicate 95% confidence regions (multivariate t-distribution). Dots represent individual samples and are colored by geographical regions. The values in parentheses on the *x* and *y* axes represent the proportion of variance explained by each principal component.

To further evaluate the efficacy of nanopore sequencing in identifying CpGs within complex and functionally significant genomic regions, we compared our findings with three WGBS datasets (GSE181854^24^, GSE186458^11^, and GSE80911^25^). Although the total number of CpGs detected by ONT and WGBS was similar, nanopore sequencing identified a higher proportion of CpGs in gene-related regions (36.9%), regulatory regions (21.9%), repeat regions (50.9%), regions near CpG islands (CGIs, 54.0%), and telomere-adjacent regions (7.4%) than in the three WGBS datasets (Figs. 2c-f). Thus, nanopore sequencing in our study provides enhanced characterization of methylation patterns, particularly in critical genomic regions.

Following the completion of high-quality, haplotype-phased methylation calling, we aggregated the CpGs from all individuals to construct an extensive methylation atlas comprising 27,609,354 CpGs. Notably, the majority (90.9%) of CpGs are commonly shared among individuals in the Chinese population, with population frequencies exceeding 0.9. Moreover, these commonly shared CpGs exhibit methylation levels surpassing 85%, with no significant variation observed in this methylation pattern across the studied populations (Fig. 2g).

To explore the genomic region-specific characteristics of the methylation atlas, we performed functional annotation of the aggregated CpGs, focusing on gene-related regions, regulatory regions (including histone marks and DNase hypersensitivity sites), CGI-related regions, repeat regions, and near telomere regions. Our analysis indicated that the vast majority of functional elements show no significant variation in CpG counts across the three studied populations. We further evaluated the density and methylation levels of CpGs within each functional element (Fig. 2h). Introns, TSS200, DNase hypersensitivity sites, low complexity regions (LCRs), retroposons, and CGIs all demonstrated a higher density of CpGs compared to their respective genomic divisions. Additionally, among these elements, TSS200, DNase hypersensitivity sites, LCRs, and CGIs were characterized by lower methylation levels, while retroposons exhibited higher methylation levels. Notably, these regions corresponded to where nanopore sequencing detected a greater number of CpGs.

To investigate potential methylation differences among populations, we conducted principal component analysis (PCA) on commonly shared CpGs that exhibited variable methylation levels. As illustrated in Figs. 2i-l, individuals from the North, South, and Xizang regions display distinct methylation patterns, with North and South partially intermingled while Xizang diverging from both. To quantify these observations, we conducted an analysis of similarity (ANOSIM), which revealed notable methylation divergence between the high-altitude population and the low-altitude inland populations (Bray-Curtis ANOSIM: Xizang vs. North = 0.72, Xizang vs. South = 0.49), whereas populations from low-altitude inland areas exhibit more similar methylation patterns (North vs South, Bray-Curtis ANOSIM=0.28).

### SV-related methylation patterns reveal genome-wide compensatory and demethylation mechanisms

SVs are a prevalent type of mutation in human genomes. Besides incurring genetic perturbations, SVs are shown in recent studies to exhibit epigenetic effects, with their methylation states implicated in the suppression of transposable elements, genome-wide epigenetic remodeling, and aging^26, 27, 28^. On a whole-genome level, we investigated the methylation patterns associated with four types of SVs: duplications (DUP), deletions (DEL), insertions (INS), and inversions (INV), while a genome-wide methylation background was constructed to account for inter-person variabilities. Among the four types of SVs, the average methylation levels of DUP and INV were within the genome background levels. Interestingly, DEL showed significantly higher methylation levels than the background, while INS exhibited lower methylation levels than the background (Fig. 3a).

**Figure 3.**
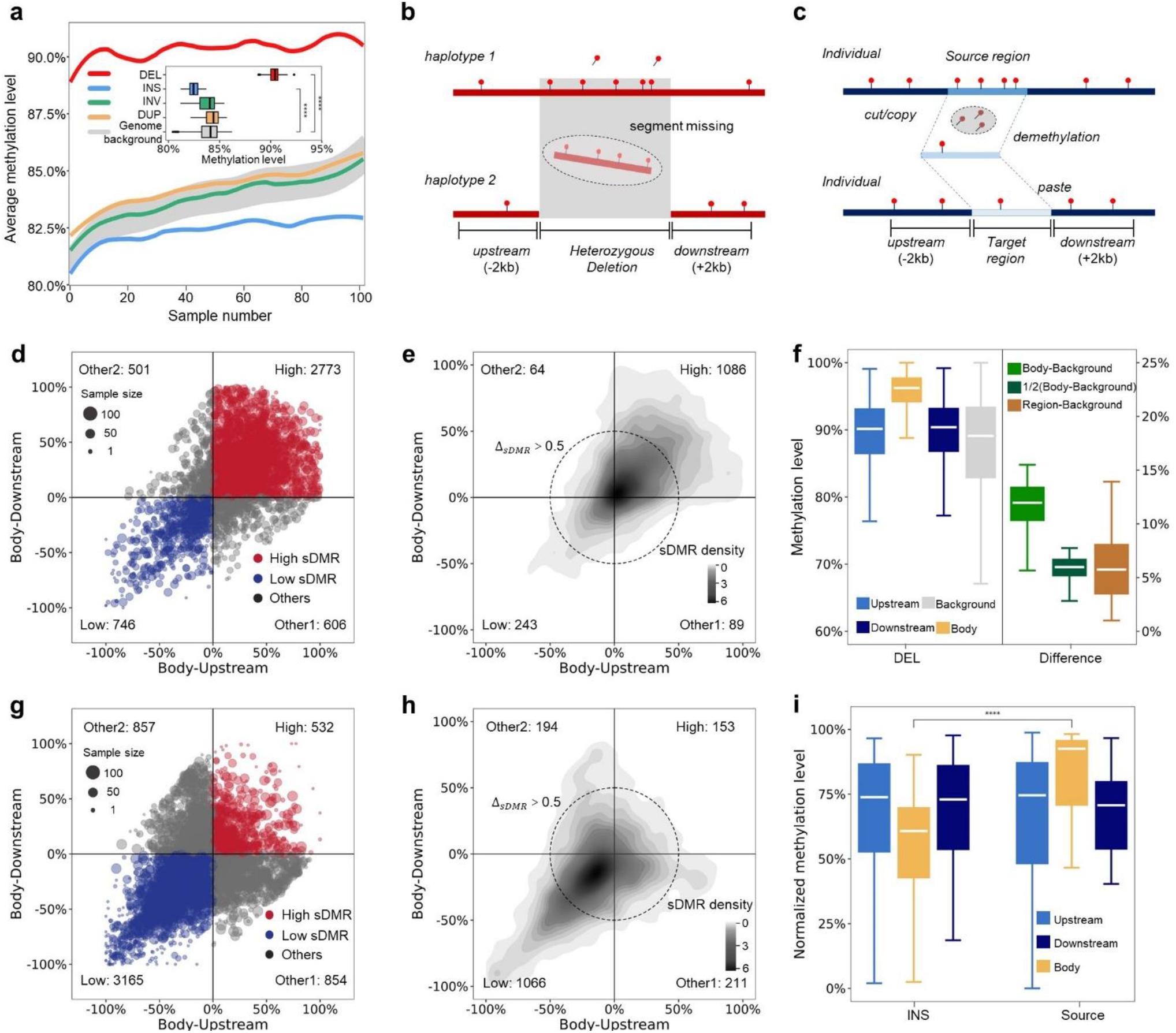
SV-associated methylation patterns identified by sDMR analysis. **(a)** Methylation level distribution on SVs for each of the 106 individuals, with gray shading delineating the range of genomic background methylation. The inset boxplot shows the methylation level distribution of each SV type aggregated across all individuals. **(b, c)** Schematic of methylation dynamics associated with deletions **(b)** and insertions **(c). (d, e)** Methylation differences **(d)** and sDMR density **(e)** for heterozygous deletions. The *x*-axis represents the normalized methylation level of the deletion body minus that of its 2 kb upstream region, while the *y*-axis shows the difference between the deletion body and its 2 kb downstream region, both calculated after normalization. In (**d**), raw sDMRs are shown and classified into four groups: High, Low, and two other categories. In (**e**), filtered sDMRs (Δ_*sDMR*_ > 0.5, dashed circle) are visualized by density. **(f)** Methylation levels in deletion bodies minus upstream/downstream regions, normalized to genomic background. Right panel shows the distribution of methylation differences between the indicated regions. **(g, h)** Methylation differences **(g)** and sDMR density **(h)** for insertions. The calculation and classifications are the same as in panels (**d)** and **(e**). **(i)** Methylation levels of MEI sites (INS) and their original source loci (Source).

This result prompted us to examine the differential methylation of DEL and INS against their associated genomic segments (Figs. 3b-c). We calculated Δ_*sDMR*_, the difference in average methylation levels within a genomic segment versus in its upstream and downstream 2k bp regions, to identify high and low sDMRs. For DELs, Δ_*sDMR*_ was calculated for the undeleted allele at all heterozygous DEL loci. The results indicated that DELs are distributed toward the “High sDMR” quadrant where the undeleted segment is hyper-methylated compared to both upstream and downstream flanking regions. Consistently, the number of hyper-methylated DELs (*n*=1,086 by Δ_*sDMR*_ > 0.5) was significantly greater than the numbers in other quadrants (Figs. 3d-e).

We further quantified the segmental methylation differences against the background methylation distribution of 1000 bp (the median DEL length) sliding windows in individual genomes. Surprisingly, we found at all loci, the difference in methylation level between the undeleted segment and its flanking regions is around 2-fold of the difference between the flanking regions and the genomic background (Fig. 3f). That is, once one allele of a segment is deleted from the diploid genome, the remaining allele displays doubled hyper- or hypo-methylation in excess to its genomic background, suggestive of a linear compensatory effect in epigenetic regulation. The compensation slightly varies for hyper-(1.94-fold) and hypo-methylated (2.3-fold) DELs, but is robust on the individual level and within the range of 250–6,500 bp of DEL length. Furthermore, the fold compensation was highly consistent across exons, introns, promoters and CGIs, regions pertinent to gene expression. These include genes involved in diverse fundamental processes, such as *CHORDC1* (molecular chaperone), *SIK3* (a key regulator of metabolic and circadian processes), and *SSUH2* (mRNA processing and transcription termination). Therefore, we speculate that the pronounced two-fold compensation at DEL loci across broad genome contexts may provide a sensitive mechanism for maintaining the normal expression dosage of genes despite potential interference by SVs. Similarly, we calculated Δ_*sDMR*_ for INS segments, and found a systematic distribution of INS toward hypomethylation with respect to flanking regions (the “Low sDMR” quadrant, *n*=1,066 by Δ_*sDMR*_>0.5, Figs. 3g-h). From these INSs, we were able to isolate 33 high frequency (population frequency>50%) mobile element insertions (MEIs) whose source locations could be reliably determined, including 2 MEIs of the cut-and-paste type and 31 MEIs of the copy-and-paste type. This allowed us to investigate the changes in methylation levels between the MEI sites and their source locations. While we found no significant methylation differences between the 2k bp flanking regions in the source versus the destination sites, the methylation levels in the MEIs were significantly lower than those in the source elements (Fig. 3i). The systematic demethylation of MEIs has been reported in cancer somatic mutations, which might have reflected transposition-associated epigenetic erasure^29^. However, for germline MEIs, we observed the alteration in methylation levels with respect to the source element scales with the population frequency of the MEI. This result suggested that the re-establishment of epigenetic suppression on mobile elements might have been counteracted by selective forces maintaining the demethylation state at the insertion site over evolutionary timescales. In fact, twelve of these MEIs were proximal to gene regions, of which six were located in exon or promoter regions. Representative genes include *CHST6, ADAMTS6*, and *GNPDA1*, critical for corneal development, cardiovascular formation, and glucose metabolism, respectively ^30, 31, 32^.

### Haplotype-based methylation patterns uncover population-specific imprinted genes

Leveraging the haplotype phasing capabilities of long-range spanning nanopore reads, we randomly selected ten individuals from each population to analyze differentially methylated regions between the paternal and maternal chromosomes at a haplotype scale (hDMRs) across individuals. The median per sample hDMR counts identified in the North, South, and Xizang populations were 273, 248, and 152, respectively, with the corresponding Δ_*hDMR*_ of 0.56, 0.55, and 0.64, respectively. For all three regions, the biallelic methylation levels on hDMRs are distributed independently of the overall genomic methylation pattern (Figs. 4a-c). However, Xizang exhibited significantly different hDMR counts (p=0.014) and methylation differences (p=0.026) when compared to the South and North. In particular, the hDMRs in the Xizang population are more densely concentrated in regions with greater methylation differences (Fig. 4c). Notably, no significant differences were observed for the higher methylation level between haplotypes among the North, South and Xizang populations (0.84, 0.829 and 0.831, respectively). In contrast, significant differences (p < 0.001) were detected in the methylation levels of the less methylated haplotype (Fig. 4d). These results suggest that the greater Δ_*hDMR*_ at hDMRs in the Xizang population is a result of pronounced methylation erasure at the de-suppressed chromosome.

**Figure 4.**
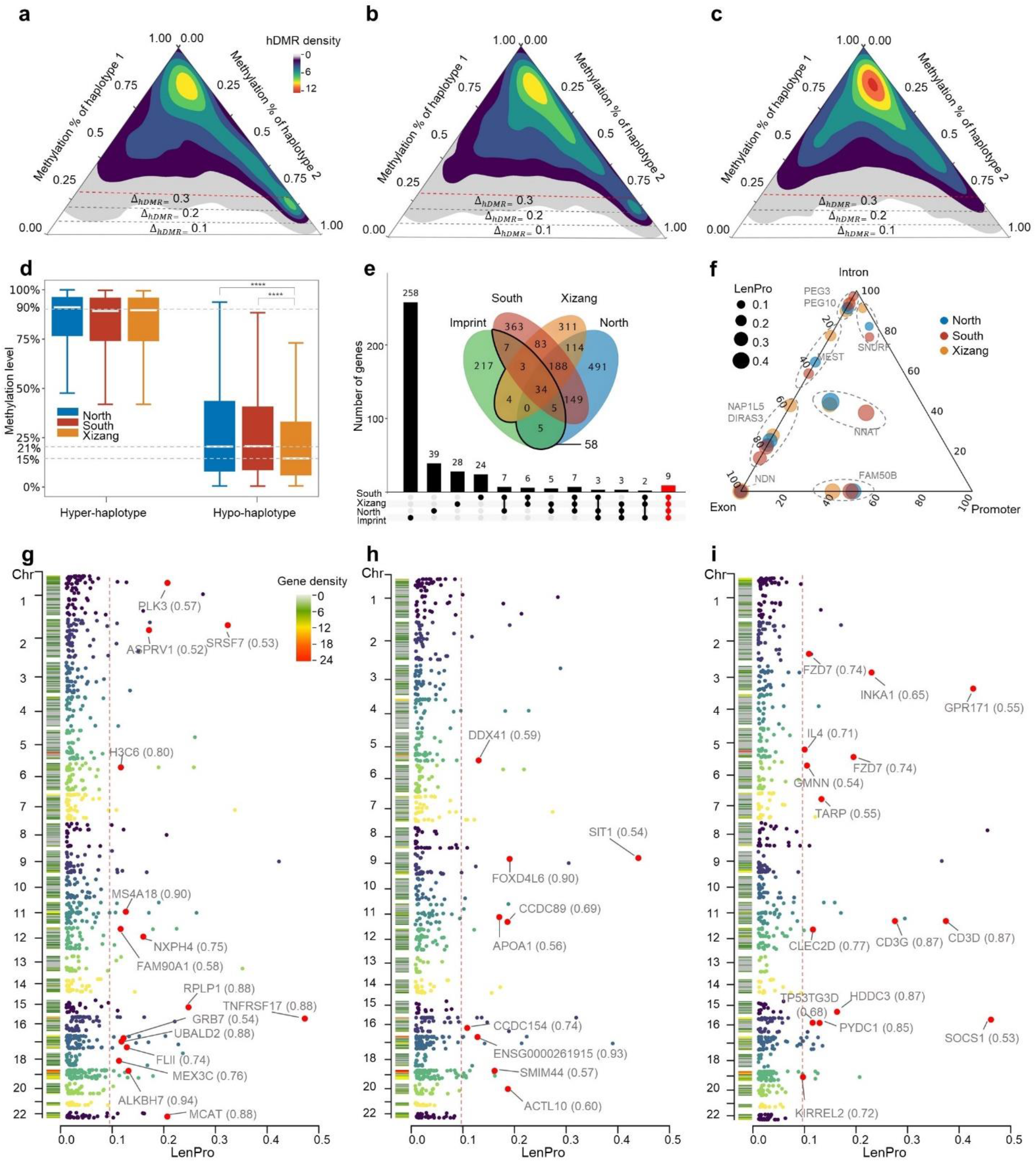
Population-specific imprinting regions identified through hDMR analysis. **(a–c)** Density distribution of biallelic methylation levels on hDMRs in **(a)** North, **(b)** South, and **(c)** Xizang populations. Dashed lines denote Δ_*hDMR*_ thresholds of 0.1, 0.2, and 0.3, with the red dashed lines highlighting the cutoff used in this study. Proximity to the upper vertex indicates stronger haplotype-specific methylation differences. **(d)** Distribution of methylation levels on the hypo- and hyper-methylated haplotypes for hDMRs in three populations. Dashed lines indicate median values: 15% for hypomethylated haplotypes in the Xizang population, 21% for the hypomethylated haplotypes in the North and South populations, and 90% for the hypermethylated haplotypes across all three populations. **(e)** Overlap between discovered hDMRs and known imprinted genes. Venn diagram shows intersections for hDMRs of Δ_*hDMR*_ > 0.3, with the black box indicating 58 genes verified by the public GENEIMPRINT database. UpSet plot displays stringent overlaps (hDMRs with Δ_*hDMR*_ > 0.3 and LenPro > 0.1), with 9 genes shared across all three populations (highlighted in red). **(f)** hDMRs associated with 9 imprinted genes shared across populations, showing the relative proportions of hDMR length overlapping exon, intron and promoter regions (*x* + *y* + *z* = 100%). Each point represents a hDMR in one population. hDMRs belonging to the same gene are enclosed with dashed circular outlines. **(g–i)** Population-specific candidate imprinted genes in Northern **(g)**, Southern **(h)**, and Xizang cohorts **(i).** High-confidence genes (Δ_*hDMR*_ > 0.5 and LenPro > 0.1) are highlighted in red dots. Gene names are annotated with their corresponding Δ_*hDMR*_ values in parentheses. The red dashed line denotes the LenPro threshold of 0.1. All other hDMRs (dark blue, green and yellow dots) are colored by their chromosome locations

To further investigate the potential impact of hDMR, we selected hDMRs with Δ_*hDMR*_> 0.3. We found that the majority (>60%) of hDMRs are shared by less than 10% of samples, showing highly individualized patterns. Further PCA analysis based on these hDMRs revealed that samples from the North and South regions overlapped to span the entire principal component space, whereas samples from Xizang displayed substantially more uniform hDMR signatures. This finding might have resulted from the variegated eco-environmental factors that the inland populations are exposed to and their higher genetic exchange rate. In contrast, the Xizang population might have been epigenetically adapted to dominating environmental factors.

Using these hDMRs, we identified a total of 1,709 candidate imprint control regions (ICRs), of which 58 were verified by the public GENEIMPRINT databases. A comprehensive comparison with previously reported ICR datasets^33, 34^ further confirmed the reliability of our identified ICRs. Upon filtering by the proportion of hDMR length to gene length (LenPro > 0.1), we identified nine imprinted genes (*DIRAS3, FAM50B, MEST, NAP1L5, NDN, NNAT, PEG10, PEG3, SNURF*) shared by three populations and confirmed by GENEIMPRINT (Fig. 4e). The ICRs within these common imprinted genes are distributed across exon, intron, and promoter regions, exhibiting no significant location preference between gene-related regions among the three populations (Fig. 4f). Meanwhile, from ICR-associated genes absent from the GENEIMPRINT database, we identified 15, 9, and 15 genes unique to the North, South, and Xizang populations, respectively (Figs. 4g-i). Notably, the genes in the North population (such as *PLK3, ASPRV1*, and *MCAT*) are involved in stress responses, skin protection, and energy metabolism regulation, which are essential for adapting to cold environments. The genes in the South population (such as *DDX41* and *APOA1*) play roles in immune regulation and metabolic processes, supporting adaptation to humid environments. Meanwhile, genes in the Xizang population (such as *GPR171, IL4*, and *LTC4S*) are associated with immune modulation, asthma, and lipogenesis, which are critical for survival at high altitudes^35, 36, 37^. Given the diversity of hDMRs between populations and individuals, we propose that they influence individuals’ environmental adaptability in a more dynamic fashion, particularly by fine-tuning haplotype-specific methylation levels within gene regions.

### Population-specific differential methylation reveals genes involved in high-altitude adaptation

Given the substantial environmental differences among the three populations, we identified population-specific pDMRs from pairwise comparisons of the North, South and Xizang populations. Our findings revealed a total of 297 pDMRs between the North and South (Fig. 5a). In contrast, the comparisons between Xizang and North, as well as Xizang and South, yielded significantly higher numbers of pDMRs (965 and 1,078, respectively, Figs. 5b-c). Besides overall pDMR counts, a higher proportion of hypo-pDMR (regions of lower methylation in the focal population) was identified between Xizang and the low-altitude inland areas. We found that genes, SINE elements, and inter-CGI regions accumulate higher numbers of pDMRs. However, after normalization by the length of functional regions, pDMRs are more densely enriched in specific regions including TSS200 (5.8%), CGIs (4.0%), SSRs (8.5%), LCRs (11.8%), and retroposons (3.0%) (Fig. 5d). Notably, with the exception of retroposons, all these areas typically exhibit lower levels of methylation. The enrichment of pDMRs in TSS200 and CGI can be considered a contribution of regulatory regions to environmental adaptability^38^. The accumulation of pDMRs in SSR, LCR, and retroposon may be associated with the high variability and flexibility of these areas, which render them sensitive targets for epigenetic regulation^39^.

**Figure 5.**
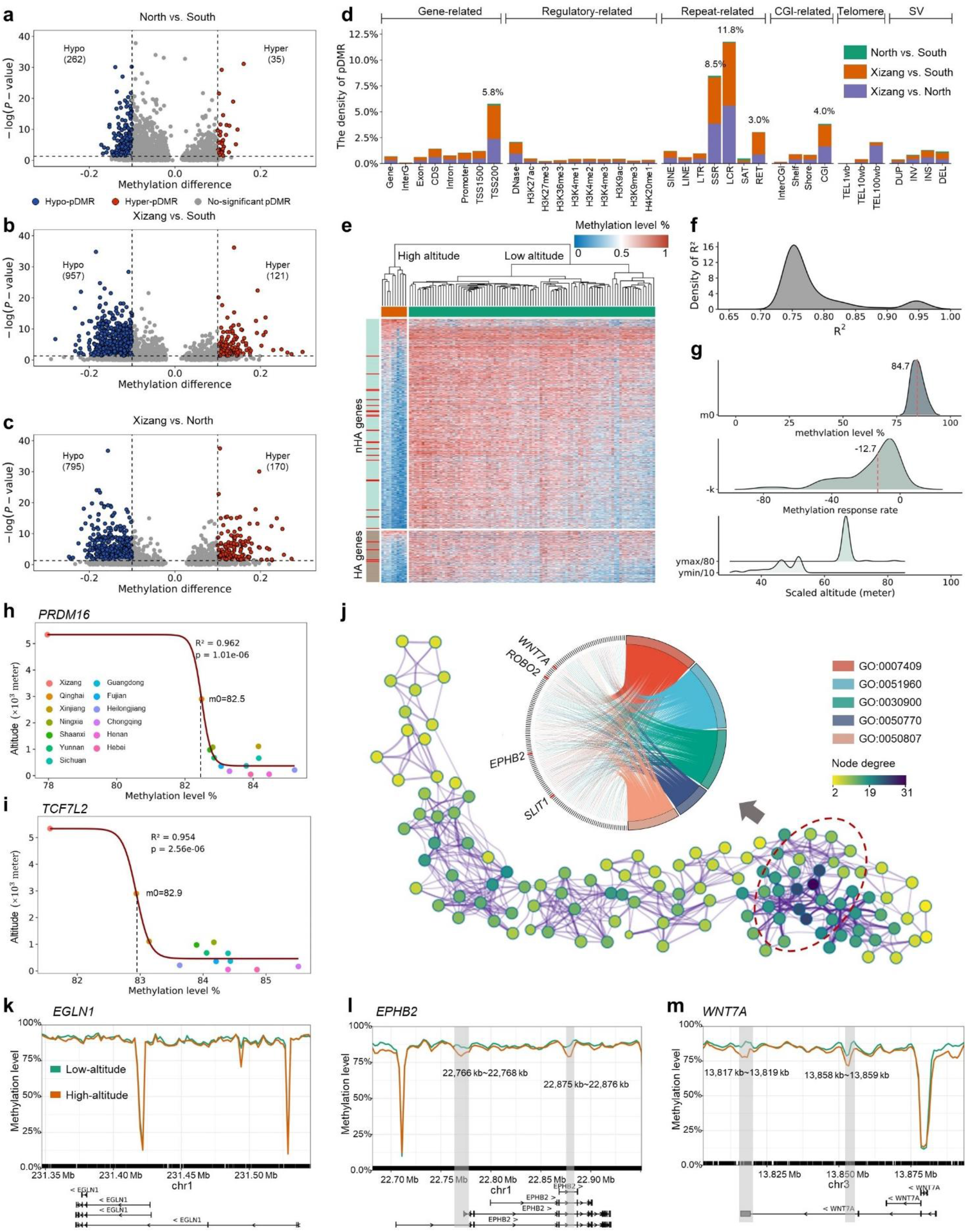
High-altitude adaptive epigenetic signatures identified through pDMRs. **(a-c)** pDMR identification for North vs. South **(a)**, Xizang vs. South **(b)**, and Xizang vs. North **(c)** populations. The numbers in parentheses indicate the counts of hypomethylated and hypermethylated pDMRs identified in each pairwise comparison. Dashed horizontal lines indicate the p-value threshold of 0.05, while dashed vertical lines represent the methylation difference threshold of ±0.1. **(d)** pDMR densities across genome functional regions. **(e)** Altitude-associated methylation patterns. The heatmap presents normalized methylation levels per gene, with columns grouped by gene type (nHA vs. HA) and rows grouped by population (low-altitude vs. high-altitude populations). Population clusters (top) were determined by hierarchical clustering. Genes highly responsive to altitude in the fitting analysis (R^2^ > 0.9 and all fitted parameters have p < 0.05) are indicated by red bars. **(f-g)** Fitted parameter distributions for altitude responsive DMGs. Parameters include: R^2^ of sigmoidal curve fitting **(f)** and m_0_, k, and altitude bounds of *y*_*min*_and *y*_*max*_**(g).** Red dashed lines show the median values for m_0_ and k. **(h-i)** Altitude-methylation response curves for candidate adaptive genes *PRDM16* **(h)** and *TCF7L2* **(i)**. Dashed lines indicate the position of m_0_, representing the inflection point in the fitted sigmoidal curve. **(j)** Network enrichment analysis color-coded by node connectivity. Red circles highlight key pathway clusters. Chord diagrams visualize genes shared by the top five pathways (indicated by red bars). **(k-m)** Methylation levels along three high-altitude adaptive genes in Xizang (high-altitude) and inland (low-altitude) populations, including a known adaptive gene *EGLN1* **(k)** and two novel adaptive candidates *EPHB2* **(l)** and *WNT7A* **(m)**. Gray shading indicates significant pDMRs; gene structures are shown below each track.

To compare the impacts of different environmental factors on genome methylation, we annotated these pDMRs and their associated differential methylated sites onto gene functional regions to identify differentially methylated genes (DMGs). For the altitude factor (Xizang vs. inland, including North and South), we identified 1,408 DMGs, of which 24 genes have been reported to associate with high-altitude adaptation^40^. For the temperature/humidity (TH) factor (North vs South), we identified 203 DMGs, of which only 6 genes are clearly related to TH regulation. We further conducted KEGG and GO enrichment analyses. With regard to altitude-associated DMGs, we selected the top 20 clusters and found HIF-1 signaling pathway (hsa04066) was significantly enriched (p-value = 1.43×10^−10^). This pathway plays a critical role in maintaining oxygen homeostasis^41^. Concurrently, we identified pathways associated with blood circulation, vascular processes, and heart regulation, which collectively contribute to the regulation of systemic processes (GO:0044057, p-value = 1.1×10^−26^). These pathways can indirectly influence adaptation to high-altitude environments^42, 43^. The enrichment results of TH-associated DMGs primarily encompass metabolic regulation, cellular function and homeostasis, and DNA repair processes, without a direct association with adaptation to TH. These findings suggest that altitude differences exert a more substantial impact on methylation levels than TH differences.

Given the pronounced differential methylation observed between Xizang and inland populations, we conducted a thorough investigation of altitude-associated DMGs. We categorized altitude-associated DMGs into two groups: publicly reported high-altitude adaptation (HA) genes (≥ 2 literature support), and potentially novel high-altitude adaptation (nHA) genes. Cluster analysis revealed divergence of Xizang and inland populations on methylation levels of both HA and nHA genes (Fig. 5e). Moreover, compared to HA genes, nHA genes exhibited a larger effect size (Cliff’s Delta = 0.398) and greater mean differences (0.043) between methylation levels in Xizang vs. inland populations. To establish a quantitative model for epigenetic high-altitude responses, we fitted, for all HA and nHA genes, the altitudes of the population habitats to the gene’s methylation levels by a sigmoidal curve. The fitting was based on the hypothesis that DNA methylation modulates gene expression by classical repressor dynamics and the gene expression dosage scales with altitude. Remarkably, almost all fittings were significant (>74.9% HA and nHA genes having R^2^> 0.7, Fig. 5f). Moreover, a subset of genes (*n* = 61) formed a minor peak at R^2^> 0.9. We suspect that these genes’ epigenetic states are dominated by high-altitude associated environmental factors. Surprisingly, the key fitting parameter *m*_0_, the methylation level inducing half of the altitude response range, tightly situated around 84.7%, and *k*, the linear response rate at *m*_0_, spanned a wide range with a median of -12.7 (Fig. 5g). Interestingly, the fitted values of *m*_0_ align with the estimated sample-wise genomic background methylation of 80.5%-86.1% (Fig. 3a). This result suggests that genome-wide methylation is set at a level of maximal tunability for epigenetic regulation, and also partially explains the linear compensation effect at heterozygous deletion sites (Fig. 3f). By requesting p-values < 0.05 for all fitted parameters, we further screened 24 genes extremely well-fitted to the sigmoid response curve (red barred in Fig. 5e, Figs. 5h-i). This gene set is predominantly composed of nHA genes such as *PRDM16* and *TCF7L2* – both were involved in fat and glucose metabolism, and their roles may be linked to metabolic regulation under high-altitude conditions^44, 45^. In contrast, genes with reduced R^2^ in the range 0.7 – 0.9 (major peak in Fig. 5f) may be pleiotropic or subject to other mechanisms of regulation.

Lastly, to uncover genes of high-altitude adaptation from a gene interaction network perspective, we constructed an enrichment network by integrating the 20 enrichment clusters mentioned previously. The five most interconnected pathways share nHA genes, such as *EPHB2, ROBO2, SLIT1*, and *WNT7A*, which are enriched in the functional module associated with nervous system development (Fig. 5j). We further compared the methylation levels of these four genes with three previously identified HA genes associated with hypoxia adaptation and lipid metabolism (*EPAS1, EGLN1*, and *PPARA*)^46, 47, 48^. Interestingly, the three HA genes do not contain significant pDMRs in Xizang vs. inland samples (Fig. 5k). In comparison, *EPHB2* and *WNT7A* showed notable differences in methylation levels in their regulatory and coding regions (Figs. 5l–m), suggesting significant epigenetic regulation. Such genes may represent epigenetically regulated network hubs contributing to the high-altitude adaptability in the Xizang population.

### An open-source methylation atlas for the Chinese population

The data presented in this study is compiled and made available on a web server ChinaMeth (http://bioinformatics.hit.edu.cn/chinaMeth/#/). ChinaMeth offers major statistics, as well as comprehensive and interactive views of the methylation atlas for Chinese populations. Users can examine the CpG density and methylation levels in continuous 1 kb windows along the genome with aligned gene annotations. One individual from each of the three populations is randomly selected for display. In addition to the methylation landscape, ChinaMeth catalogues sDMR, hDMR, and pDMRs identified in this study. sDMRs are displayed with the normalized methylation levels of the structural variation itself and the surrounding 2 kb regions. For hDMRs, users can view the position, length, and methylation levels for ten individuals from each population. The pDMR section displays all detected pDMRs without filtering, offering comprehensive comparisons across populations. To enhance user accessibility, we provide two search options: by gene name and by chromosomal location. This online platform serves as a valuable tool for understanding methylation patterns in the Chinese population and facilitating case investigations into the impact of epigenetic regulation on population characteristics.

## Discussion

In this study, we constructed a comprehensive methylation atlas of the Chinese population, based on data from 106 healthy individuals inhabiting a wide range of geographical areas in the North, South, and Xizang regions. This unprecedented methylome collection encompasses 27,609,354 CpGs genome-wide, and captures a spectrum of methylation modifications, from those commonly shared to those uniquely personal, offering new avenues for investigating methylation patterns within the Chinese population. In comparison to previous WGBS-based studies, the use of nanopore long-read sequencing not only offers enhanced characterization of methylation profiles in gene-related regions, but also allows the analysis of methylation patterns with respect to large SVs and megabase-scale haplotypes. Paralleling the adoption of long-read sequencing in cataloging methylomes across tissues and developmental stages^11^, this study presents the largest population-scale DNA methylation atlas of Chinese individuals generated using Nanopore sequencing, offering high-resolution, whole-genome methylation data with broad population representation. In conjunction with the diverse habitats for samples of a controlled genetic ancestry, reveals epigenetic mechanisms and epigenetic adaptation on evolutionary timescales.

In the interrogation of DNA methylation with respect to genomic structural variants, we observed a previously unreported phenomenon – heterozygous deletions are epigenetically compensated to roughly 2-fold of its surrounding regions. While it is tempting to interpret this phenomenon as a response to haploid loss of epigenetic regulation, whether this *cis-* type of epigenetic compensation translates into gene dosage compensation is elusive. The classic epigenetic dosage compensation, e.g. in X-chromosome inactivation, acts in *trans*, i.e., one copy of the diploid genes is fully methylated to abolish ½ of diploid gene expression. However, for a 2-fold epigenetic compensation in *cis* to restore the associated gene expression dosage, a linear dose-response model must hold for gene expression regulation by DNA methylation. Although experimental evidence is lacking, reassuringly, our quantitative analysis of pDMRs across a wide altitude range indicated that, at least for altitude-responsive genes, all their fitted dose-response curves collapsed on one inflection point (*m*_0_) coinciding with the genomic background methylation level. This result suggested that it could be a general principle for the genome background methylation to fall into a range of linear regulatory response, such that *cis*-acting epigenetic compensation within its vicinity could be utilized to finetune gene expression against DNA fragmental loss. The 2-fold epigenetic compensation is consistently observed across various lengths or contexts of the deleted fragments. Additionally, we analyzed the epigenetic compensation effect in independent samples of different age groups. We observed a steady trend of decay in the compensation fold among the young, middle aged and old groups. This trend suggests the epigenetic compensation to haploid DNA segment loss may be driven by an active mechanism that degrades with the aging process. Therefore, an interesting question to pursue is the underlying molecular mechanism for haploid loss detection and epigenetic compensation.

In addition, we observed global demethylation on insertions, especially on MEIs by directly comparing methylation on its source and on the inserted element. Previous studies proposed demethylation as a mechanism of transposable element mobilization^26, 49^, and suggested that transposition itself could be prone to demethylation enzymes^50^ or activate DNA damage repair associated epigenetic remodeling^51^. These processes pertain to the short-term dynamics of transposable element epigenetics. In comparison, the observed demethylation on a population scale offers a fresh perspective that these MEIs could be selectively de-repressed for the epigenetic regulation of proximal gene expression. The consequences of such global MEI de-repression and whether it promotes future transposition warrant further in-depth exploration.

Through the population-scale hDMR and pDMR analyses, we observed distinctive patterns associated with the Xizang population against inland populations. Xizang possesses a lower number of hDMRs, possibly reflecting a mixed effect of stable population inheritance with low external genetic exchange, and phenotypic adaptation to relatively controlled environmental factors. However, for both hDMR and pDMRs, the Xizang population exhibits more pronounced methylation differences through pervasive demethylation, mostly in SSR and LCRs. These are likely regions providing the genome with epigenetic regulatory flexibility. Based on pDMRs, we found that altitude exerts substantial impacts on the apparent DNA methylation in Chinese populations compared to other environmental factors. Leveraging the altitude gradient of the selected populations, we established quantitative response functions for DMGs. This analysis revealed a gene set whose epigenetic state is dominated by altitude-associated environmental factors. Combined with analysis from the pathway and network perspectives, we found that known high-altitude adaptation genes like *EGLN1, EPAS1*^15^ were not regulated epigenetically, whereas newly identified hub genes such as *EPHB2* and *WNT7A* exhibit epigenetic hallmarks in their regulatory regions in the Xizang population. Our stringent analysis pipeline and multi-faceted interrogation substantially expanded the potential set of high-altitude adaptation genes^52, 53^ and population specific imprinted genes, providing a rich genomics resource for further studies on molecular and functional linkages of epigenetically controlled phenotypic adaptation. It is worth noting that genetic divergence such as SNVs may govern methylation at specific CpG sites^18^. Given the complex genetic backgrounds of the populations being studied, we evaluated the potential effects of methylation quantitative trait loci (meQTL) on the identification of hDMRs and pDMRs. To this end, we integrated our previously published SNV panel of the Chinese population in the China100K project^54^ with public meQTL resources^55^, and found that SNVs affect at most 2-7% of pDMRs and hDMRs identified in this study, although this represents a highly conservative overestimation. Therefore, the observed methylation differences are unlikely to be driven by genetic variation, but rather reflect independent epigenetic processes.

It is important to note that the methylation modifications reported in the current atlas primarily involve 5mC. The inclusion of other DNA modifications, such as 5hmC and 6mA, will greatly enhance the power of the atlas to facilitate research in epigenome dynamics and fine-grained epigenome regulation^56, 57^. However, compared to the identification of 5mC, greater effort is required to improve the relatively lower performance of the current calling approaches for 5hmC and 6mA. On the other hand, although the current atlas successfully resolved methylation modifications across nearly all genomic regions, particularly in regions difficult for WGBS, certain regions of potential functional significance—such as the rDNA array, large-scale segmental duplications, and centromeres—remain inadequately characterized^58, 59^. Therefore, our future objective is to establish a comprehensive methylation discovery framework at the telomere-to-telomere level, enabling the detection of all types of methylation across any genomic region.

## Acknowledgements

We thank the Center for Bioinformatics at the Harbin Institute of Technology for providing the sequencing and data analysis platform that supported this work.

## Data availability

The SVs data generated by ONT sequencing are available from the Genome Variation Map (GVM, https://ngdc.cncb.ac.cn/gvm/) at the National Genomics Data Center, Beijing Institute of Genomics, Chinese Academy of Sciences, and the China National Center for Bioinformation, under accession number GVM000420. The reference SNV dataset representing allele frequency differences in the Chinese population is available under accession number GVM000418. The cohort CpG data have been submitted to the Open Archive for Miscellaneous Data (OMIX, http://ngdc.cncb.ac.cn/omix/) under accession number OMIX006130. The three types of DMRs identified during our study, along with an interactive web portal for data visualization, can be accessed at http://bioinformatics.hit.edu.cn/chinaMeth/#/. The three WGBS datasets used in this study can be downloaded using GEO accession numbers GSE181854, GSE186458, and GSE80911 from NCBI.

## Code availability

The plotting and processing scripts is available at https://github.com/YLeeHIT/ChinaMethAtlas and is distributed under MIT License.

